# High-efficiency transformation and gene expression in *Picosynechococcus* sp. PCC 7002

**DOI:** 10.1101/2024.09.17.613521

**Authors:** Andrew P. Hren, Joshua P. Abraham, Melissa P. Tumen-Velasquez, Michael Melesse Vergara, Adam M. Guss, William G. Alexander, Brian F. Pfleger, Jerome M. Fox, Carrie A. Eckert

## Abstract

Cyanobacteria are promising microbial platforms for a myriad of biotechnological applications, from sustainable biomaterials to photosynthetic chemical production, but still lack the breadth of genetic tools available for more commonly engineered microbes such as *Escherichia coli*. This study develops genetic tools to enhance the transformation efficiency and heterologous gene expression in *Picosynechococcus* sp. PCC 7002, a fast-growing, halotolerant, and naturally competent strain. Integration of fluorescent reporter cassettes across the genome revealed an integration site that yields a fourfold improvement in gene expression relative to previously reported sites. Protocol optimization and engineered DNA methylation in *E. coli* increased transformation efficiency by over tenfold. This work provides an experimental framework for efficient genome editing and metabolic engineering in the model cyanobacterium PCC 7002.

## Introduction

As the sole bacterial phylum capable of oxygenic photosynthesis, cyanobacteria such as *Picosynechococcus* sp. PCC 7002 (also known as ATCC 27264, hereafter PCC 7002) are of growing interest to scientists and metabolic engineers. Isolated from Magueyes Island in Puerto Rico (Gotto et al., 1979), PCC 7002 is tolerant to high light intensities (Nomura et al., 2006) and high salt concentrations (Batterton & Van Baalen, 1971). It is a promising host for converting wastewater and CO_2_ streams directly to value-added products (Korosh et al., 2018) or for creating microbial biomass that can be used for heterotrophic conversion of renewable feedstocks to bioproducts (Comer et al., 2020; Hays & Ducat, 2015).

PCC 7002 is naturally transformable (Frigaard et al., 2004) and can integrate DNA from plasmids or linear fragments into its genome via homologous recombination (Xu et al., 2010), a capability exploited in several, established protocols (Dallo et al., 2022; Ruffing et al., 2016; Xu et al., 2010). PCC 7002 also has an annotated genome (RefSeq: GCF_000019485.1) and a wide variety of advanced methods which provide a foundation for rational strain engineering (Begemann et al., 2013; Gordon et al., 2016; Hendry et al., 2016; Markley et al., 2015; Perez et al., 2017; Ruffing et al., 2016; Ungerer & Pakrasi, 2016; Zess et al., 2016). However, PCC 7002 still lacks some techniques in its toolkit, such as a list of validated neutral insertion sites for genomic integration. As the level and consistency of genomically-integrated, heterologous genes expression is locus dependent, putative integration loci require direct experimental validation to quantify potential expression levels (Bryant et al., 2014; Chaves et al., 2020). While a handful of studies have identified expression-neutral loci within the PCC 7002 genome (Ruffing et al., 2016; Vogel et al., 2017; M. Wang et al., 2019), the relative expression levels of foreign genes within these sites remains unexplored.

Another shortcoming of the current PCC 7002 toolkit is the lack of a native restriction-modification (R-M) system workaround. R-M systems are prokaryotic immune systems that protect bacteria against selfish genetic elements (Vasu & Nagaraja, 2013). Methylation patterns introduced by methyltransferases serve as a self/non-self recognition system that enables a cell to detect and degrade foreign DNA (Zhao et al., 2006). These R-M systems pose a major barrier to genetic transformation in many microorganisms, though their effects can be countered by methylating a DNA payload via heterologous expression of R-M methyltransferases in a cloning strain of *E. coli* (Riley & Guss, 2021). Expression of methyltransferases native to *Synechocystis sp*. PCC 6803, a well-studied freshwater cyanobacterium, increased transformation efficiency of harvested DNA up to two orders of magnitude in *Synechocystis* (B. Wang et al., 2015) and in PCC 7002 (Kamoku & Nielsen, 2024). Production of similar cloning strains using native PCC 7002 methyltransferases requires methylome analysis of the cyanobacterium, highlighting a third shortcoming of the existing PCC 7002 toolkit.

In this study, we sought to expand the genetic engineering toolkit for PCC 7002 by characterizing additional integration loci, standardizing a high efficiency natural transformation protocol, and enhancing the transformation efficiency via pre-methylation of recombinant DNA. These standardized protocols will facilitate future research in PCC 7002 by enabling high throughput, library-scale techniques.

## Results and Discussion

### Characterization of Chromosomal Loci for Gene Expression in PCC 7002

In prior work, the expression of an integrated enhanced yellow fluorescent protein (EYFP) gene was used as a reporter to characterize engineering-relevant loci and synthetic promoters in PCC 7002 (Begemann et al., 2013; Markley et al., 2015). Using a similar approach, we investigated several loci with physiological relevance and compared them against the established Δ*acsA* locus (Begemann et al., 2013), an acrylic acid based counter-selectable marker. Constructs containing EYFP reporter under IPTG induction (P_cLac143_) were introduced into wildtype PCC 7002 by natural transformation and selected with CRISPR-Cas12a to ensure marker-less integration. Maximum IPTG-induced fluorescence varied across sites, with both higher- and lower-expressing loci relative to Δ*acsA* (Figure 1A). Notably, the highest expression occurred with integration of the EYFP cassette into the native plasmid pAQ3. Prior work suggests that, under normal salinity conditions and early growth stages, the copy number of pAQ3 is three times that of the chromosome (Xu et al., 2010). Elevated copy number is a strong indicator of total gene expression *in vivo* (Chandler & Pritchard, 1975) and likely contributes to the high observed fluorescence. For chromosomal loci, the highest fluorescence was observed upon integration downstream of *cpcF*, an essential gene of the phycocyanin pathway. Prior studies have used the *cpc* operon promoter (P_cpc_) as a starting point for high constitutive expression (Markley et al., 2015), and these data support the high relative expression afforded by the *cpc* operon, relative to other loci on the PCC 7002 chromosome. Interestingly, EYFP expression from neutral site 1 (NS1), between genes A0932 and A0933, was greater than expression of EYFP downstream of essential proteins, such as RuBisCO (*rbcLXS*) and the D subunit of photosystem II (*psbA*). These data suggest that proximity to highly expressed genes does not necessarily yield high heterologous expression of a reporter gene.

**Figure 1.**
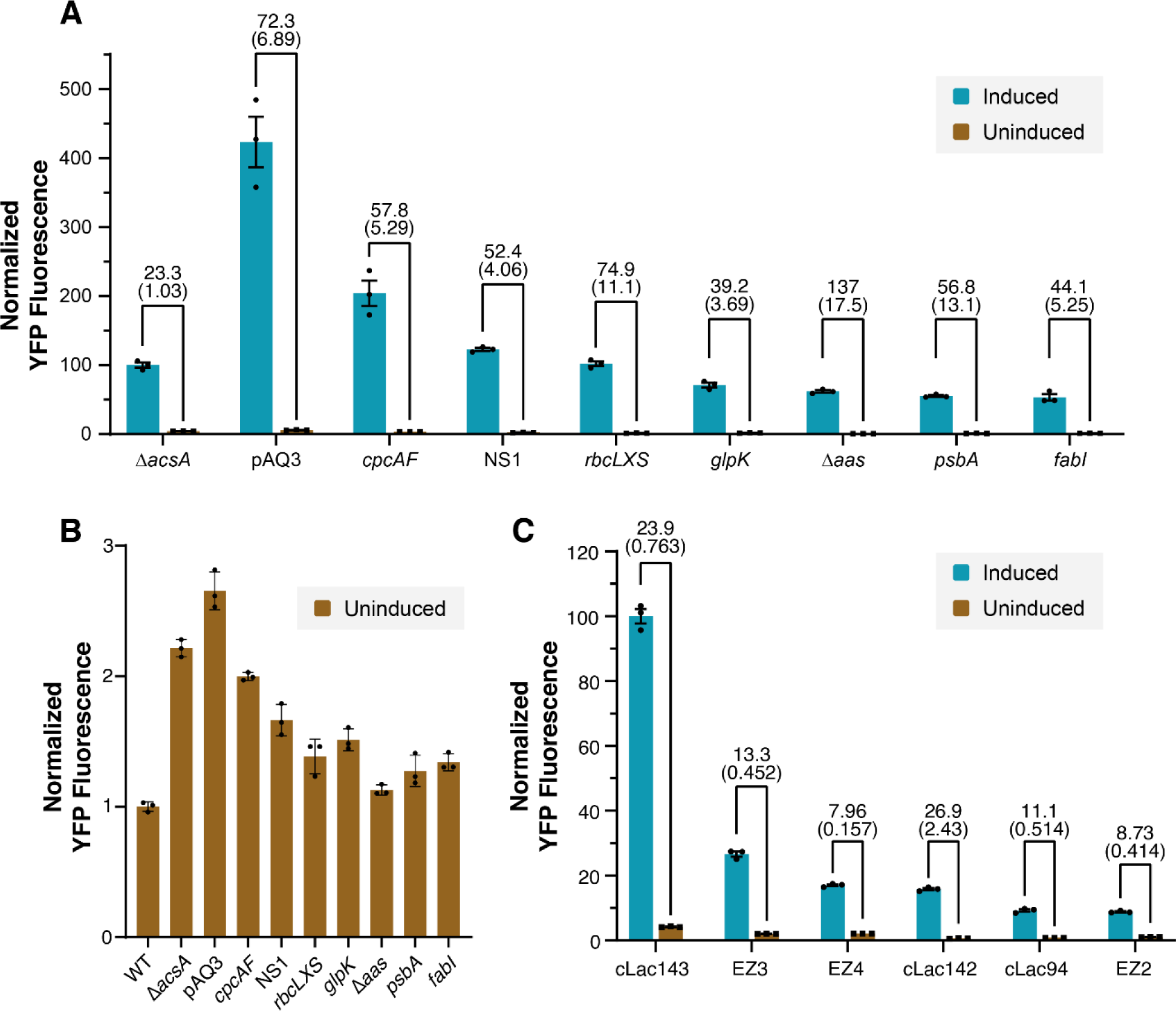
Relative expression of EYFP across integration sites and inducible promoters. (A) Relative EYFP fluorescence of induced and uninduced EYFP reporter cassettes integrated into 9 different loci, normalized to mean induced fluorescence of *ΔacsA* with the P_cLac143_ promoter. Enumerated data above each bar cluster indicates the fold-change in EYFP fluorescence in each locus and its standard error. (B) Fluorescence of only the uninduced states normalized to wildtype PCC 7002. (C) Relative EYFP fluorescence from six unique promoters fused to the EYFP reporter cassette integrated into *ΔacsA*. cLac promoters were induced at 1 mM IPTG and EZ promoters at 1000 ng mL^-1^ aTc. Error bars indicate standard error of the mean for n=3 biological replicates. The numbers above the bars represent the fold difference between induced and uninduced samples, with the corresponding standard error in parentheses.

The dynamic range (i.e., difference in expression between induced and uninduced states) is a valuable metric for the level of control of gene expression afforded by an inducible promoter. We observed a high dynamic range (137-fold) for the Δ*aas* locus, a relevant target for engineering fatty acid biosynthesis (Kaczmarzyk & Fulda, 2010), much higher than that observed for the Δ*acsA* locus (23-fold). When comparing the relative fluorescence magnitudes of the uninduced states across loci, we observed a high level of leaky expression at Δ*acsA* (Figure 1B). Due to the large magnitude of fluorescence in the uninduced state relative to its induced state, leaky expression has an outsized influence on dynamic range in this locus. We explored how alternative inducible promoters (Zess et al., 2016) behave within the Δ*acsA* locus and found that P_cLac143_ exhibited higher basal fluorescence and also the highest maximal induced fluorescence, with a high dynamic range (Figure 1C), consistent with previous work (Markley et al., 2015). Promoter P_cLac142_ also showed a high dynamic range, with substantially lower basal expression and lower maximal expression than P_cLac143._

Disparate growth across uninduced strains suggests that some tested loci have deleterious fitness effects. Cassettes integrated downstream of *fabI*, *psbA* and *glpK* and in place of *acsA* caused strains to grow significantly worse in the first 24 hours compared to wildtype (Supplementary Figure 2A; statistical significance determined using a Tukey HSD test). The growth defects observed in *glpK* and Δ*acsA,* common integration sites in prior work (Begemann et al., 2013; Gordon et al., 2016; Hren et al., 2024; Selão et al., 2019; Vogel et al., 2017), highlight the importance of direct experimental characterization of integration sites that are often assumed to be neutral. As fitness effect, expression level, and dynamic range of inducible systems are strongly influenced by a chosen integration site, our data on the locus-dependent behavior of heterologous cassettes can inform future genetic engineering efforts in PCC 7002.

### Transformation Protocol Optimization

High-throughput methods (e.g., genome-scale CRISPR screens) require high-efficiency transformation protocols to ensure maximal coverage and minimal bias of large libraries. Unfortunately, the efficiencies of published PCC 7002 transformation protocols vary widely due to a range of unoptimized conditions (Begemann et al., 2013; Essich et al., 1990; Frigaard et al., 2004; Gupta et al., 2020; Kachel & Mack, 2020; Ruffing, 2014; Selão et al., 2019; Stevens & Porter, 1980; Vogel et al., 2017; Xu et al., 2010). To begin, we tested the effect of cell density, homology arm length, and recovery time on the transformation efficiency of a test plasmid, pAPH07 (Hren et al., 2024), which contains a kanamycin resistance marker and a gRNA for CRISPR/dCas9 flanked on either side by homology arms that direct insertion downstream of *glpK*. We found that longer homology arms (i.e. increasing from 750 bp to 1250 bp on each side) increased efficiency of integration into the *glpK* locus by 3.4-fold (Figure 2A), a phenomenon observed in previous work (Ruffing et al., 2016). Cells (i) harvested at OD_730_ of 0.3 and (ii) concentrated to an OD_730_ of at least 3.0 enhanced efficiency as a function of cell density (Figure 2B). In turn, increasing incubation time (i.e., 1 to 24 h) enhanced efficiency further (Fig. 2C). When combined, the adjustments in protocol enhanced the transformation efficiency of pAPH07 by an order of magnitude (i.e., 1.36 x 10^5^ to 1.58 x 10^6^) over a previously described protocol (Fig. 2D) (Ruffing et al., 2016). We anticipate that this method for efficient integration will facilitate preliminary library-scale experiments and high-throughput studies of PCC 7002, with additional improvements necessary at elevated library sizes.

**Figure 2.**
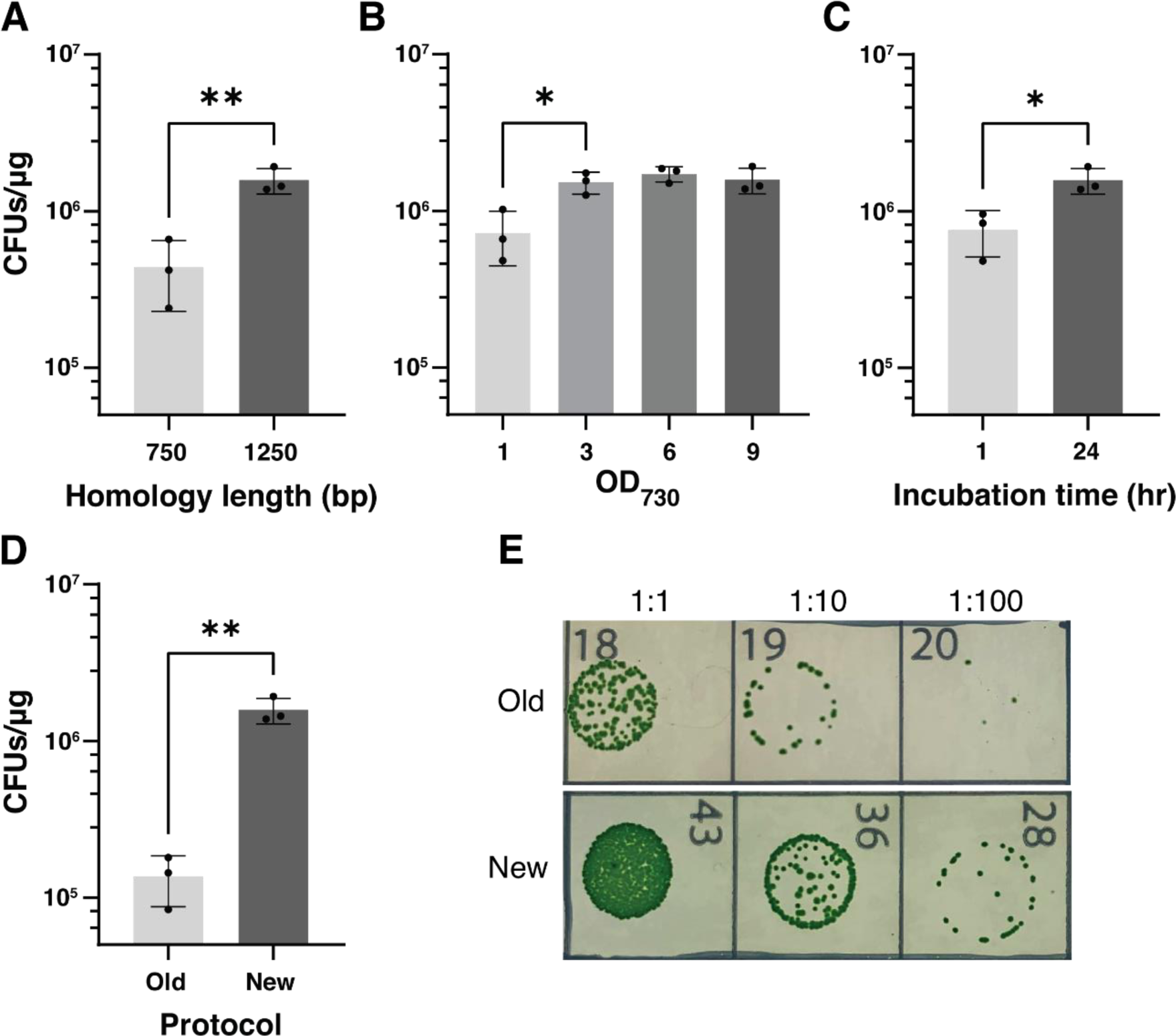
Transformation protocol optimization. (A-C) The transformation protocol of pAPH07 (Hren et al., 2024) into PCC 7002 was improved by (A) lengthening vector homology arms, (B) increasing starting cell density during transformation, and (C) increasing the length of incubation. Variables were set to default levels when not directly tested (1250 bp, OD_730_=9.0, 24 hours) (D) The updated protocol showed an order of magnitude improvement over the initial protocol adapted from previous work (Ruffing et al., 2016). (E) Images of serial dilutions used to calculate transformation efficiency, with the new protocol generating more colony forming units (CFUs). Data depicts n=3 biological replicates per condition.

### Methylome Analysis Identifies Methylated Motifs in PCC 7002

We first examined the predicted methyltransferases encoded in the PCC 7002 chromosome using the New England Biolab’s Restriction Enzyme Database, or REBASE (Roberts et al., 2023), which lists predicted methylation sites and methyltransferase-encoding genes based on comparisons to known methyltransferases in related organisms. As R-M systems are highly variable between strains of bacteria, we sought to validate these predictions using the newly-developed methylome discovery software package MIJAMP (Tidwell et al., 2024), which relies on Oxford Nanopore Technology’s sequencing systems to call modified bases and discover the motifs in which they are embedded. MIJAMP also allows for direct querying of the data gathered for specific motifs, allowing us to interrogate the methylation status of potentially missed motifs.

Our results (Table 1) largely agree with those predicted by REBASE with some differences. The activity of one enzyme (M.AquORF12P) predicted to methylate the YGGCCR motif was not identified during motif discovery. The evidence for this motif in REBASE, however, appears shallow, as the motif listed does not include the precise location of the 5mC modification. Direct query of this motif in the dataset confirmed no methylation at either cysteine, refuting the REBASE prediction. Our analysis also suggests an alternative methylation pattern for M.AquORF849P: G(5mC)GATCGC exhibited a genome-wide methylation (GWM) score of almost 99% (i.e. ∼99% of those motifs were methylated in this dataset), while the predicted (5mC)GATCG from REBASE only exhibited ∼92% GWM. Finally, we detected three additional methylation motifs (TGG(6mA)GG, CCC(4mC)GC, and GG(6mA)N_5_CTTC) not assigned to a methyltransferase by REBASE.

**Table 1:** Methylation Motifs in PCC 7002.

| Motif | Methyltransferase<br>(REBASE) | Methylated Sites | Total Sites | Percent<br>Methylated |
| --- | --- | --- | --- | --- |
| G(6mA)TC | M.AquDam (A1620) | 44886 | 45782 | **98.04** |
| G(5mC)GATCGC<sup>a</sup> | M.AquORF849P<br>(A0849) | 11040 | 11170 | **98.84** |
| TGG(6mA)GG |  | 1784 | 1831 | **97.43** |
| CCC(4mC)GC |  | 1528 | 1800 | **84.89** |
| GAGG(6mA)G | AquIII (A2132) | 1398 | 1453 | **96.21** |
| (5mC)YCGRG | M.AquI (A1188 –<br>A1188-2) | 1217 | 1448 | **84.05** |
| GRGGA(6mA)G | AquIV (A2314) | 1259 | 1271 | **99.06** |
| GCCGN(6mA)C | AquII (A0358) | 1218 | 1230 | **99.02** |
| GA(6mA)GN<sub>5</sub>TCC |  | 334 | 340 | **98.24** |
| GG(6mA)N<sub>5</sub>CTTC |  | 337 | 340 | **99.12** |
| CRA(6mA)N<sub>8</sub>TGAC | M.AquV (C0003-C0004) | 259 | 260 | **99.62** |
| GTC(6mA)N<sub>8</sub>TTYG | M.AquV (C0003-C0004) | 244 | 260 | **93.85** |

<sup>a</sup>REBASE predicts this motif to be (5mC)GATCG

### Expression of DNA Methyltransferases in *E. coli* Increases PCC 7002 Transformation Efficiency

To determine if DNA methylation would protect plasmid DNA from restriction during transformation in PCC 7002, we generated strains of *E. coli* (AG11039 and AG11304, Table 2) that are deficient in native DNA methyltransferases and instead express PCC 7002 methyltransferases whose methylation motifs were detected in our methylome analysis and are associated with restriction enzymes (Table 1; additional strains are listed in Supplementary Table 5). We confirmed the activity of the heterologous methyltransferases via nanopore sequencing and subsequent analysis using MIJAMP (Table 3). In both strains, we detected the expected methylation motifs for all methyltransferases except for A0358 (GCCGN(6mA)C), potentially due to poor expression or incompatibility in the *E. coli* host. Similarly, the lower GWM score of the (5mC)YCGRG motif in AG11304 (∼77%) could be attributed to low expression or enzyme activity; however, this same motif also exhibited lower GWM scores (∼84%) in the PCC 7002 methylome analysis. These data highlight the importance of experimentally validating methylation targets when expressing DNA methyltransferases in *E. coli*.

**Table 2:**
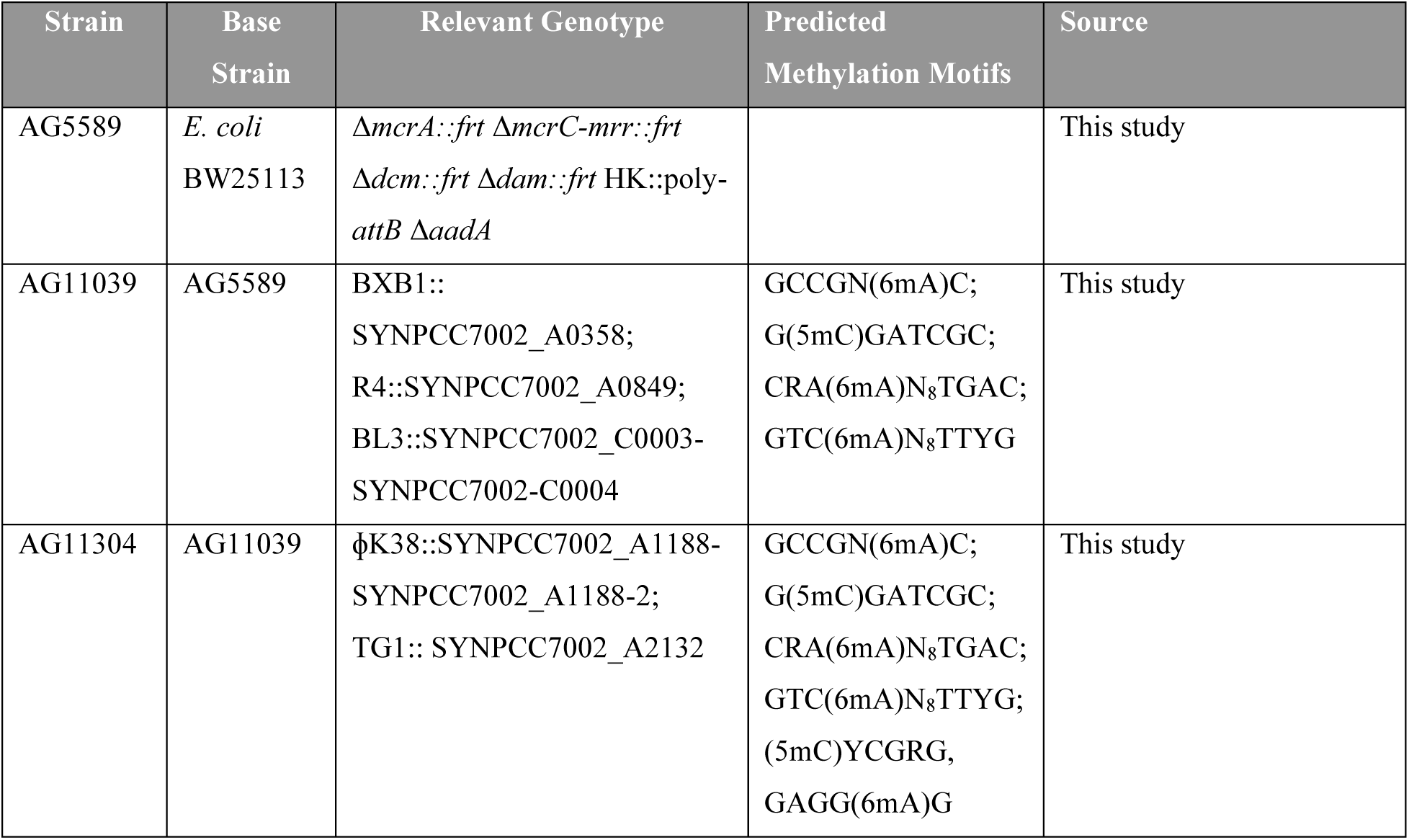
*E. coli* Methyltransferase Expression Strains.

**Table 3:** Identified Methylation Motifs in *E. coli* Shuttle Strains.

| Strain | Motif | Methylated Sites | Total Sites | Percent Methylated |
| --- | --- | --- | --- | --- |
| AG11039 | G(5mC)GATCGC | 426 | 440 | **96.82** |
|  | CRA(6mA)N<sub>8</sub>TGAC | 368 | 374 | **98.40** |
|  | GTC(6mA)N<sub>8</sub>TTYG | 347 | 374 | **93.85** |
| AG11304 | (5mC)YCGRG | 1920 | 2482 | **77.36** |
|  | GAGG(6mA)G | 724 | 737 | **98.24** |
|  | G(5mC)GATCGC | 436 | 440 | **99.09** |
|  | CRA(6mA)N<sub>8</sub>TGAC | 372 | 374 | **99.47** |
|  | GTC(6mA)N<sub>8</sub>TTYG | 357 | 374 | **95.45** |

We transformed these methylation strains, along with DH5α, with several test plasmids (pJA167, pJA168, pJA173, and pJA204) containing a gentamicin resistance cassette flanked by 750 bp of homology directed to several integration sites (Δ*acsA*, *glpK*, *fabI*, and Δ*lexA*). We transformed the control and the methylated DNA into wildtype PCC 7002, performed serial dilutions, and measured transformation efficiency. In each tested plasmid, the three methyltransferases expressed in AG11039 significantly increased the transformation efficiency (Figure 3A-D). Interestingly, improvement to transformation efficiency varied between the tested plasmids. Alignment of the motifs from Table 3 on the tested cassettes (Figure 3E) provides some insight to the observed trends. For example, while the Δ*acsA* locus experienced less than a threefold improvement using either AG11039 or AG11304, integration downstream of *glpK* had improvements in efficiency nearly 30-fold. We hypothesize that the presence of the CRA(6mA)N_8_TGAC motif in the downstream *glpK* homology region contributes to the drastic increase in efficiency when methylated. Moreover, the extra methylation from AG11304 only further enhanced transformation efficiency of the cassette inserted downstream of *fabI*. This increase is supported by the presence of the (5mC)YCGRG motif only in the *fabI* homology arm, compared to the others tested. These data suggest that the observed improvements to transformation efficiency are dependent on the presence and methylation of motifs detected by our analyses. In all cases, methylation from either AG11039 or AG11304 significantly enhanced transformation efficiency relative to plasmids derived from *E. coli* DH5α.

**Figure 3.**
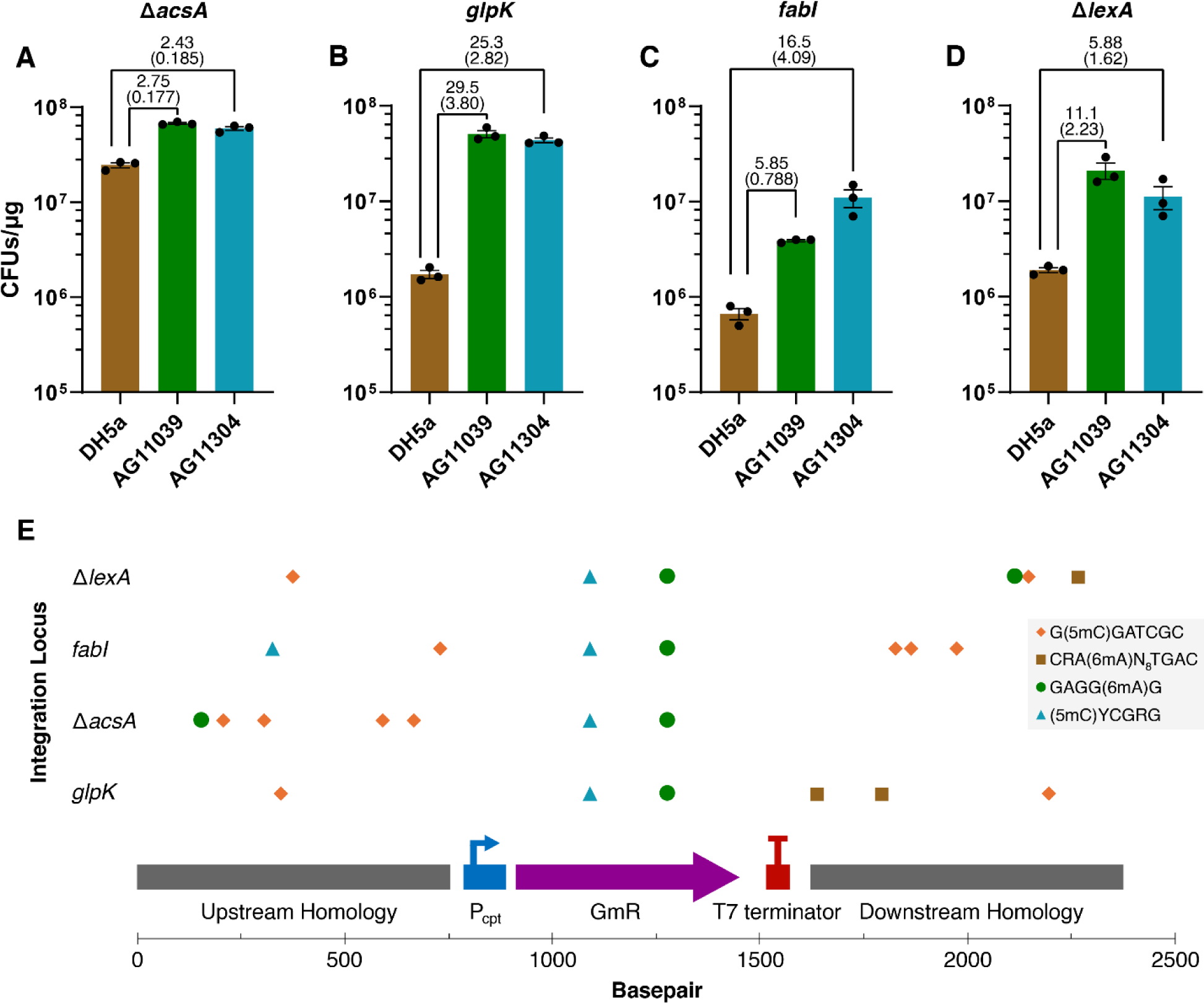
Methylation of DNA with PCC 7002 methyltransferases dramatically improves transformation efficiency. Wild-type PCC 7002 was transformed with a gentamicin resistance cassette integrating into (A) Δ*acsA,* (B) *glpK*, (C) fabI, and (D) Δ*lexA*. All strains exhibited significant increases in transformation efficiency when plasmids were methylated, which was calculated using an unpaired Student’s t-test (p<0.0001). (E) Location of methylated motifs from AG11039 and AG11304 (Table 3) are shown for each cassette tested in A-D.

## Conclusions

In this work, we characterized the locus-dependent behavior of heterologous expression cassettes in *Picosynechococcus* sp. PCC 7002. Additionally, we present an improved protocol for high efficiency natural transformation and two strains of *E. coli* that express methyltransferases from PCC 7002. Plasmids purified from these strains demonstrate an order of magnitude improvement in transformation efficiency compared to controls. These methylation strains, in conjunction with our recommended transformation protocol, can provide researchers with opportunities to generate large libraries or difficult mutants of PCC 7002. We anticipate that these additions to the PCC 7002 toolkit will provide a base for future work towards high-efficiency genetic manipulation.

## Materials and Methods

### Cyanobacterial Culturing

PCC 7002 was grown from frozen strains stored at -80°C in Media A+ (Supplementary Table 7) with a cryoprotectant, either (20% (v/v) glycerol or 10% (v/v) DMSO). Experiments were initiated from single colonies from freezer stocks isolated on Media A+ plates (1% agar). Cells were grown in three formats: 20 mL glass bubble tubes (foam stoppers, ambient air sparging, and 50 µmol photons m^−2^ s^−1^ full-spectrum LED light) for overnight culturing; 250 mL baffled shake flasks (plastic caps, ISF1-X Kuhner shaker at 160 RPM, 1% CO_2_, and 250 µmol photons m^−2^ s^−1^ full-spectrum LED light) for fluorescence experiments; and 125 mL Erlenmeyer flasks (foam stoppers, 200 RPM, air, 150 µmol photons m^−2^ s^−1^ full-spectrum LED light) for transformation efficiency experiments. Antibiotics were added to working concentrations of 50 µg mL^-1^ kanamycin or 35 µg mL^-1^ gentamicin, depending on genetic cassette integrated.

### Standardized Transformation for PCC 7002

Wild-type PCC 7002 was inoculated into 25 mL Media A+ (initial OD_730_ = 0.05; Thermo Scientific Genesys 10S Vis) and grown overnight in the Erlenmeyer flask format until an OD_730_ of 0.2-0.3 was reached. Cultures were centrifuged at 4,300 rcf for 10 minutes, the supernatant was decanted, and then pellets were resuspended in Media A+ to an OD_730_ of 1.0-9.0 (optimal = 9.0). In a 1.5 mL microcentrifuge tube, 300 µL of concentrated cells were combined with 0.1 – 1 µg of plasmid DNA (quantified by Thermo Scientific NanoDrop) and vortexed to mix, then incubated at 37°C and 150 µmol photon m^-2^ s^-1^ for 1-24 hours (optimal = 24). The tube was vortexed briefly to resuspend settled cells, then plated directly onto selective Media A+ plates or serially diluted with Media A+ and spotted (5 µL) to quantify transformants. Plates were incubated at 37°C with ambient air and white light (100–150 µmol photon m^-2^ s^-1^) until colonies were visible. Transformation efficiency *η* was calculated as the number of colony forming units (CFUs) per µg of DNA, incorporating spot volume *V_s_*, dilution factor *d*, total transformation volume *V_t_*, and mass of DNA added *m,* as described in Equation 1.

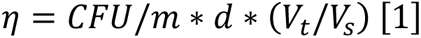

For efficiency comparisons, we tested a modified version of a published transformation protocol (Ruffing et al., 2016). Briefly, PCC 7002 was grown to an OD_730_ of 1.0 in the Erlenmeyer flask format, then 1 mL was transferred to a 16 mm glass test tube with a foam stopper and mixed with 100 ng of linearized plasmid DNA (pAPH07; NdeI digestion). The transformation was incubated for 24 hours (200 RPM, 30°C, 60 µmol photon m^-2^ s^-1^), then efficiency was calculated via spot plating.

### Construction of Repair Plasmids

A modular Golden Gate Assembly-based workflow was established for quick generation of constructs that could be introduced into a characterized locus. We ordered 750 bp homology arms up and downstream of the target integration site as synthesized plasmids from Twist Biosciences. These homology arms contained a “Golden Gate Adapter” that could be cleaved with BsmBI and ligated with fragments corresponding to the desired promoter, gene of interest, and terminator/repressor. These fragments were obtained by PCR (NEB Q5 High-Fidelity DNA Polymerase) to introduce BsmBI restriction sites with appropriate overlaps. An example 20 µL Golden Gate Assembly would contain a 2:1 molar ratio of each insert to the homology vector (typically 50 ng) with 1 µL Esp3I, 0.2 µL T4 DNA ligase, and 2 µL of 10X T4 Ligase Buffer. The reaction was performed in a thermocycler, with 60 cycles of digestion (37°C, 5 minutes) and ligation (16°C, 10 minutes), followed by a final digestion (37°C, 10 minutes) and heat inactivation (65°C, 20 minutes). The assembled product was then transformed into chemically competent *E. coli* DH5α for selection.

### CRISPR-Cas12a-mediated Editing of PCC 7002

pDC011 variants with gRNAs that target the loci (Supplementary Table 2) were generated by Golden Gate assembly using AarI and T4 DNA Ligase (NEB). pDC011 is a derivative of pSL2680 (Ungerer & Pakrasi, 2016) that instead uses an IPTG inducible promoter (P_cLac143_) (Markley et al., 2015) for increased Cas12a expression. Plasmids described in Supplementary Table 1 were used as repair templates for marker-less CRISPR-Cas12a (also known as Cpf1) editing (Ungerer & Pakrasi, 2016) and incubated with PCC 7002 using the standardized transformation protocol. Before plating, the PCC 7002/repair DNA mixture was combined 1:1 with cultures of *E. coli* HB101 harboring pDC011 derivatives and pRL443 (Elhai et al., 1997) for conjugation of the CRISPR-Cas12a machinery and gRNA cassette. The cultures of conjugal *E. coli* were washed at least three times with antibiotic-free LB medium before mixing with PCC 7002. The mixture was then plated on transferable, sterile Millipore HATF membranes on Media A+ plates with 5% LB and incubated overnight at 37°C under 150 µmol photon m^-2^ s^-1^ white light. The next day, membranes were transferred to Media A+ plates with 100 µg mL^-1^ kanamycin and 1 mM IPTG to activate expression of Cas12a and the gRNA cassette. Within 5 to 10 days, colonies appeared and were picked for colony PCR to screen for recombination of the repair template into the correct locus using GoTaq DNA polymerase (Promega) with an extended, initial denaturation step of 95°C for 10 minutes (primers listed in supplementary table 3). Successful colonies were passaged in antibiotic-free, liquid Media A+ until loss of the pDC011 plasmid, verified as loss of kanamycin resistance.

### Fluorescence Assay of YFP Reporter

Cultures of PCC 7002 were inoculated in 50 mL Media A+ in 250 mL baffled shake flasks at an OD_730_ of 0.1 and incubated for 24 hours at 37°C, 1% CO_2_, and 250 µmol photon m^-2^ s^-1^ white light. Cultures were then subject to processing and spectrophotometric measurement as reported previously (Markley et al., 2015); briefly, cultures were normalized to 2 OD_730 ·_ mL and centrifuged. Pellets were resuspended in 300 µL BugBuster Protein Extraction Reagent (Novagen), rocked at room temperature for 30 min, and centrifuged. Supernatant fluorescence was measured with an excitation of 514 nm and an emission of 527 nm with a Tecan M1000 plate reader.

### Genome Assembly and Methylome Analysis of PCC 7002

gDNA was extracted using the Quick-DNA Fungal/Bacterial Miniprep Kit (Zymo Research, Irvine, CA) and then prepared for sequencing using the Native Barcoding Sequencing Kit 24 V14 (SQK-NBD114.24); manufacturer’s instructions were followed with the exception of an omitted bead cleanup between end repair and barcode ligation. The library was loaded onto an R10.4.1 MinION flowcell, generating 1.83M reads with an N50 of 8.3 kbp. Live basecalling was performed within MinKnow using an HP Z8 workstation equipped with two NVIDIA RTX A6000 GPUs. Reads were filtered using Filtlong (https://github.com/rrwick/Filtlong) to remove reads under 1 kbp in length and to return the top 1 Gbp worth of reads by quality score. Genome assembly was conducted on this 1 Gbp read set using the Trycycler v0.5.4 workflow (Wick et al., 2021) together with Raven v1.5.3 (Vaser & Šikić, 2021), Miniasm/Minimap v0.3 (Wick & Holt, 2019), Flye v2.9.4 (Kolmogorov et al., 2019), and Medaka v1.5 (https://github.com/nanoporetech/medaka). The resulting genome is 3.4 Mbp in size and consists of six circular molecules: one chromosome, four megaplasmids, and one small plasmid.

MIJAMP (Tidwell et al., 2024) was used to detect methylated motifs within the dataset. Briefly, the 1 Gbp read set was further reduced with Filtlong to a final size of 350 Mbp, or ∼100x coverage of the genome. Read IDs from the resulting FASTQ file were extracted via the readIDExtract.py script included in MIJAMP, then the specified read IDs were used to re-basecall the dataset via Dorado v0.7.1+80da5f5 (https://github.com/nanoporetech/dorado) with v5 all-context modified base models for 6mA, 5mC, and 4mC modifications. Following the MIJAMP workflow, the resulting BAM file and previously assembled genome were preprocessed (preprocess command), then motifs were detected (motif command), revealing seven 6mA, two 5mC, and one 4mC modified motifs, with genome-wide methylation of each motif ranging from 85-99% (Table 1). The reported proportion of explained modified bases (diagnostic command) was 99.9%, 81.9%, and 80.3% for 6mA, 5mC, and 4mC modified bases, respectively.

### Construction of *E. coli* Methylating Shuttle Strains

Genes encoding predicted DNA methyltransferases from *Picosynechococcus* sp. PCC 7002 were codon optimized to *E. coli* K12 (IDT DNA) and cloned in the non-replicating integration plasmid backbone pMTV210 by GenScript (Nanjing, China). The plasmid pMTV210 was also constructed by GenScript (Nanjing, China). In this backbone, the methyltransferases genes are expressed from an arabinose-inducible promoter in *E. coli*. The Serine-integrase Assisted Genome Engineering (SAGE) method (Elmore et al., 2023) was used to integrate all the *Picosynechococcus* methyltransferases. First, pMTV1133 was integrated in AG5589. This base strain, AG5589, was engineered from AG4277 (Riley et al., 2023) by removal of the *aadA* gene using Lambda Red recombineering (Murphy & Campellone, 2003). A temperature sensitive plasmid expressing the BL3 integrase (pLAR051) was used to aid integration. 200 ng of each plasmid was transformed in 70 µL of electrocompetent cells. Transformation was recovered in 1 mL SOC at 30°C for 40 minutes to allow replication of pLAR051 and moved to 42°C for another 40 minutes for plasmid curing. The transformation was plated onto LB with 30 µg mL^-1^ kanamycin at 37°C. At least 6 colonies were picked and grown in 2 mL LB with 30 µg mL^-1^ kanamycin at 42°C overnight. Colony PCR with primers oMTV2234/oMTV2235 and oMTV24/oMTV27 were used to confirm integration of SYNPCC7002_C0003-SYNPCC7002-C0004 and the kanamycin marker in AG5589.

To remove the kanamycin marker, competent cells were made from the confirmed integration strain. Another temperature sensitive plasmid expressing the PhiC31 integrase (pLAR047) was transformed in 50 µL of competent cells and recovered in SOC at 30°C. Transformation was plated on LB with 100 µg mL carbenicillin^-1^. Colonies were picked into LB and grown at 42°C overnight. About 2 µL of the outgrowth was streaked on LB and grown overnight at 37°C. Colonies were then patched onto LB with 30 µg mL^-1^ kanamycin, LB with 100 µg mL^-1^ carbenicillin, and LB only to screen for the removal of the kanamycin resistance marker and loss of helper plasmid pLAR051. Colony PCR was used to confirm SYNPCC7002_C0003-SYNPCC7002-C0004 and the loss of the kanamycin marker to generate strain AG10460. Electrocompetent cells from AG10460 were made to integrate SYNPCC7002_A0849 (pMTV1128) using the procedure described above with pLAR058 as the recombinase helper plasmid resulting in AG10965. Similarly, SYNPCC7002_A0358 (pMTV1127) was integrated into AG10460 with pLAR074 as the helper recombinase to generate AG11039. To generate AG11078, SYNPCC7002_A2132 (pMTV1131) with helper plasmid pLAR060 was integrated in AG11039. For AG11304, SYNPCC7002_A1188-SYNPCC7002_A1188-2 (pMTV1130) was integrated using pLAR056 to aid integration.

### Methylome Analysis of *E. coli* Strains

Methylating strains were grown in 5 mL of LB with 1 mM arabinose to induce methyltransferase expression followed by high molecular weight genomic DNA extraction. Library preparation and nanopore sequencing were performed as before with *Picosynecococcus* to produce 340 Mb and 422 Mb of reads for AG11039 and AG11304, respectively. As these datasets were under 100x coverage for the *E. coli* genome, the entire read pool for each strain was used in the analysis. MIJAMP was used as with *Picosynecococcus* before to confirm methylation capability of the installed methyltransferases, however methylome discovery by MIJAMP failed to discover multiple motifs. Instead of discovering methylation *de novo*, we used the query function to check for each motif present in *Picosynecococcus* in case an enzyme’s blamed motif was incorrect (Table 3).

### *In vivo* Plasmid Methylation and Transformation of PCC 7002

Strains of modified *E. coli* BW25113 (Table 2) were transformed with PCC 7002 homology plasmids via electroporation. Briefly, cells were grown from freezer stock in LB overnight, then spun down to pellet. The supernatant was removed, then cells were resuspended in one half the culture volume of sucrose solution (300 mM) to wash. After two additional wash steps, pelleted cells were resuspended in 1/50^th^ of the culture volume of sucrose solution. Concentrated cells were electroporated at 1.8 kV in 0.2 cm cuvettes at a 1:1 DNA (ng) to cell (µL) ratio and plated on selective plates (LB agar; 35 µg mL^-1^ gentamicin). Transformed strains were grown overnight in 5 mL LB, induced at 1 mM arabinose to express the integrated methyltransferases. Methylated plasmids were purified using an Omega EZNA Plasmid Mini Kit I and used in the Standardized Transformation protocol for PCC 7002.

## Acknowledgements

The authors would like to thank Dylan Courtney for his work developing pDC011 from the original, published CRISPR/Cas12a system (Ungerer & Pakrasi, 2016).

## Data Sharing, Data Availability, and Data Reporting

The data that support the findings of this study and relevant DNA sequences are available from the corresponding author (Dr. Carrie Eckert) upon reasonable request.

## Authorship

A.H., J.A., J.F., C.E., and B.P. conceived of research and design of experiments. J.A. and A.H. constructed and tested heterologous cassettes in PCC 7002. M.T. constructed *E. coli* methyltransferase expression strains. M.T., W.G., M.V., A.G., and A.H. performed the methylome analyses. J.A. and A.H. wrote the manuscript with feedback from other authors.

