## Supplemental Materials for "High-efficiency transformation and gene expression in *Picosynechococcus* sp. PCC 7002"

Figures S1-S2. . . . . . . . . . . . . . . . . . . . . . . . . . . . . . . . . . . . . . . . . S2

Tables S1-S. . . . . . . . . . . . . . . . . . . . . . . . . . . . . . . . . . . . . . . . . S4

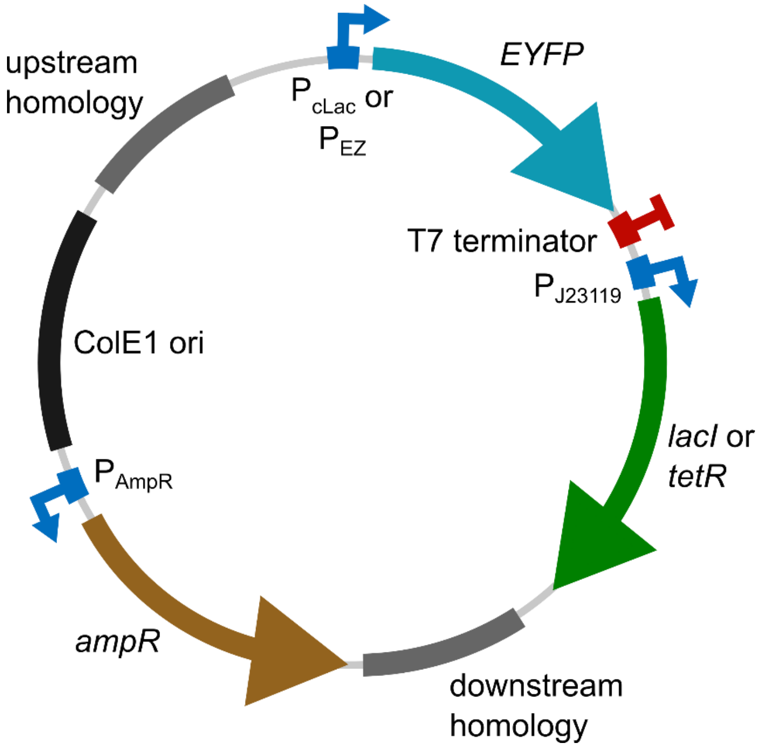


**Supplementary Figure 1. Plasmid Map for Integration of EYFP in PCC 7002.** Homologous recombination was performed using a plasmid containing 750 bp homology arms to specific loci on the PCC 7002 genome. The integrated cassettes contained a codon-optimized EYFP expressed under a corresponding inducible promoter and insulated by a T7 terminator and accompanied by a constitutive expression of the *lacI* or *tetR* repressor.

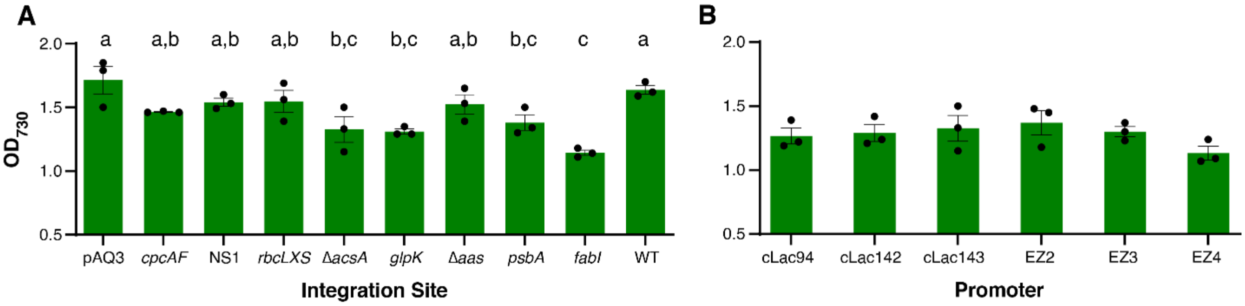


**Supplementary Figure 2. Growth effects of YFP cassette integration.** Optical density ($OD_{730}$) at 24 h measured as a proxy for cell density of PCC 7002 cultures in the (A) uninduced, locus-dependent YFP strains and the (B) uninduced, promoter-dependent strains in the Δ*acsA* locus. Significance groups are calculated based on an ANOVA and pairwise Tukey HSD test ($Q_{crit}$ > 3.948). Data depicts n=3 biological replicates per condition.

**Supplementary Table 1:** Plasmids for PCC 7002 Cloning and Testing

| Name | Relevant characteristics | Source |
|---|---|---|
| pSL2680 | RSF1010 replicating vector for PCC 7002 expression of Cas12a. Contains a *lacZ* cloning site for AarI-mediated Golden Gate assembly of gRNA, driven my $P_{J23119}$ | Ungerer 2016 |
| pDC011 | Derivative of pSL2680 that uses $P_{cLac143}$ to express Cas12a | This study |
| pRL443 | Shuttle vector for mobilizing plasmid conjugation to PCC 7002 | Elhai 1997 |
| pJA038 | Derivative of pDC011 with gRNA targeting pAQ3 | This study |
| pJA040 | Derivative of pDC011 with gRNA targeting NS1 | This study |
| pJA044 | Derivative of pDC011 with gRNA targeting Δ*acsA* | This study |
| pJA052 | Derivative of pDC011 with gRNA targeting downstream of *glpK* | This study |
| pJA054 | Derivative of pDC011 with gRNA targeting downstream of *rbcS* | This study |
| pJA057 | Derivative of pDC011 with gRNA targeting downstream of *psbA* | This study |
| pJA058 | Derivative of pDC011 with gRNA targeting downstream of *cpcF* | This study |
| pJA062 | Derivative of pDC011 with gRNA targeting Δ*aas* | This study |
| pJA100 | Derivative of pDC011 with gRNA targeting downstream of *fabI* | This study |
| pJA022 | pAQ3::$P_{cLac143}$-YFP-$P_{MB2}$-*lacI* | This study |
| pJA023 | NS1::$P_{cLac143}$-YFP-$P_{MB2}$-*lacI* | This study |
| pJA025 | Δ*acsA*::$P_{cLac143}$-YFP-$P_{MB2}$-*lacI* | This study |
| pJA029 | *glpK*::$P_{cLac143}$-YFP-$P_{MB2}$-*lacI* | This study |
| pJA030 | *rbcLXS*::$P_{cLac143}$-YFP-$P_{MB2}$-*lacI* | This study |
| pJA031 | *psbA*::$P_{cLac143}$-YFP-$P_{MB2}$-*lacI* | This study |
| pJA032 | *cpcAF*::$P_{cLac143}$-YFP-$P_{MB2}$-*lacI* | This study |
| pJA033 | Δ*aas*::$P_{cLac143}$-YFP-$P_{MB2}$-*lacI* | This study |
| pJA103 | *fabI*::$P_{cLac143}$-YFP-$P_{MB2}$-*lacI* | This study |
| pJA167 | Δ*acsA*::$P_{cpt}$-*gmR* | This study |
| pJA168 | *glpK*::$P_{cpt}$-*gmR* | This study |
| pJA173 | *fabI*::$P_{cpt}$-*gmR* | This study |
| pJA204 | Δ*lexA*::$P_{cpt}$-*gmR* | This study |
| pAPH07 | *glpK* (1250bp)::$P_{MB2}$-*lacI*+$P_{cLac143}$-sgRNA-*kanR*. Empty cassette for cloning of a CRISPRi/dCas9 guide RNA integrated downstream of *glpK* | (Hren et al., 2024) |

**Supplementary Table 2:** Guide RNA Primers for pDC011 Targeting

| Description | Sequence (5' to 3') |
|---|---|
| pAQ3_forward | AGATaacgagaaagagttatgaca |
| pAQ3_reverse | AGACtgtcataactctttctcgtt |
| *cpcAF*_forward | AGATcagcgcgggcttcttaaatc |
| *cpcAF*_reverse | AGACgatttaagaagcccgcgctg |
| NS1_forward | AGATtcagtttcatgaagcttctt |
| NS1_reverse | AGACaagaagcttcatgaaactga |
| *rbcLXS*_forward | AGATcgtctaaaattagtcgaaat |
| *rbcLXS*_reverse | AGACatttcgactaattttagacg |
| Δ*acsA*_forward | AGATctgacctgcggctaggtttc |
| Δ*acsA*_reverse | AGACgaaacctagccgcaggtcag |

| | |
|---|---|
| *glpK*_forward | AGATaactccctctaacgtaaacc |
| *glpK*_reverse | AGACggtttacgttagagggagtt |
| Δ*aas*_forward | AGATcgcttttaaatggaattgcc |
| Δ*aas*_reverse | AGACggcaattccatttaaaagcg |
| *psbA*_forward | AGATctcagcagcatctagctaaa |
| *psbA*_reverse | AGACtttagctagatgctgctgag |
| *fabI*_forward | AGATggttccccctcagtgatagg |
| *fabI*_reverse | AGACcctatcactgagggggaacc |

**Supplementary Table 3:** Primers for Verification of PCC 7002 Locus-Targeting Strains

| Description | Sequence (5' to 3') |
|---|---|
| pAQ3_seq_forward | atttcgtcagagcctacgag |
| pAQ3_seq_reverse | gcctcagtgcatatctacac |
| *cpcAF*_seq_forward | aagcctatgccgaaaatagc |
| *cpcAF*_seq_reverse | aagcgttttcgcagctttag |
| NS1_seq_forward | agacttcaccaacggttgag |
| NS1_seq_reverse | gcaccgtcttgaatggtacc |
| *rbcLXS*_seq_forward | gcgtctaattagtcagcagc |
| *rbcLXS*_seq_reverse | tccgttgccatgatgagatg |
| Δ*acsA*_seq_forward | ccgtaacctcctaggattgg |
| Δ*acsA*_seq_reverse | ctcaacccagaactacagcg |
| *glpK*_seq_forward | ctgaaaatctgtggtgccag |
| *glpK*_seq_reverse | tccgatcaccgccatctttg |
| Δ*aas*_seq_forward | caattttcgggctaggttcg |
| Δ*aas*_seq_reverse | acaatcttcttggtcagggc |
| *psbA*_seq_forward | caaggttccttctctgatgg |
| *psbA*_seq_reverse | gatttgcagctcgattatcg |
| *fabI*_seq_forward | ttctggctacgaaattatgg |
| *fabI*_seq_reverse | ccaactcttctttcaatacc |

**Supplementary Table 4**: Descriptions of all *E. coli* BW25113 Strains Generated that Express Methyltransferases from PCC 7002. Blue text represents the new methylated motif in the strain.

| Name | Relevant characteristics | Source |
|---|---|---|
| AG4277 | *E. coli* BW25113 Δ*mcrA::frt* Δ*mcrC-mrr::frt* Δ*dcm::frt* Δ*dam::frt* HK::poly-*attB* | (Riley et al., 2023) |
| AG5589 | *E. coli* BW25113 Δ*mcrA::frt* Δ*mcrC-mrr::frt* Δ*dcm::frt* Δ*dam::frt* HK::poly-*attB* Δ*aadA* | This study |
| AG10460 | BL3:: SYNPCC7002_C0003-SYNPCC7002-C0004 integrated in *E. coli* AG5589 for CRAANNNNNNNNTGAC motif methylation. Genotype: BW25113 Δ*mcrA::frt* Δ*mcrC-mrr::frt* Δ*dcm::frt* Δ*dam::frt* HK::poly-*attB* Δ*aadA* BL3:: SYNPCC7002_C0003-SYNPCC7002-C0004 | This study |
| AG10965 | R4:: SYNPCC7002_A0849 integrated in *E. coli* AG10460 for GCGATCGG and CRAANNNNNNNNTGAC motif methylation. | This study |

| | | |
|---|---|---|
| | Genotype: BW25113 Δ*mcrA::frt* Δ*mcrC-mrr::frt* Δ*dcm::frt* Δ*dam::frt* HK::poly-*attB* Δ*aadA* BL3:: SYNPCC7002_C0003-SYNPCC7002-C0004 R4:: SYNPCC7002_A0849 | |
| AG11039 | BXB1:: SYNPCC7002_A0358 integrated in *E. coli* AG10965 for GCCGAAC, GCGATCGC, and CRAANNNNNNNNTGAC motif methylation<br>Genotype: BW25113 Δ*mcrA::frt* Δ*mcrC-mrr::frt* Δ*dcm::frt* Δ*dam::frt* HK::poly-*attB* Δ*aadA* BL3:: SYNPCC7002_C0003-SYNPCC7002-C0004 R4:: SYNPCC7002_A0849 BXB1:: SYNPCC7002_A0358 | This study |
| AG11078 | TG1:: SYNPCC7002_A2132 integrated in *E. coli* AG11039 for GAGGAG, GCCGAAC, GCGATCGC, and CRAANNNNNNNNTGAC motif methylation<br>Genotype: BW25113 Δ*mcrA::frt* Δ*mcrC-mrr::frt* Δ*dcm::frt* Δ*dam::frt* HK::poly-*attB* Δ*aadA* BL3:: SYNPCC7002_C0003-SYNPCC7002-C0004 R4:: SYNPCC7002_A0849 BXB1:: SYNPCC7002_A0358 TG1:: SYNPCC7002_A2132 | This study |
| AG11304 | ϕK38:: SYNPCC7002_A1188-SYNPCC7002_A1188-2 integrated in *E. coli* AG11078 for CYCGRG, GAGGAG, GCCGAAC, GCGATCGC, and CRAANNNNNNNNTGAC motif methylation<br>Genotype: BW25113 Δ*mcrA::frt* Δ*mcrC-mrr::frt* Δ*dcm::frt* Δ*dam::frt* HK::poly-*attB* Δ*aadA* BL3:: SYNPCC7002_C0003-SYNPCC7002-C0004 R4:: SYNPCC7002_A0849 BXB1:: SYNPCC7002_A0358 TG1:: SYNPCC7002_A2132 | This study |

**Supplementary Table 5:** Plasmids used during Construction of *E. coli* Methylation Strains

| Name | Relevant characteristics | Source |
|---|---|---|
| pLAR047 | $Ap^R$; Temperature sensitive; $P_{ENO}$-jC31 recombinase | (Riley et al., 2023) |
| pLAR051 | $Ap^R$; Temperature sensitive; $P_{ENO}$-BL3 recombinase | (Riley et al., 2023) |
| pLAR053 | $Ap^R$; Temperature sensitive; $P_{ENO}$-A118 recombinase | (Riley et al., 2023) |
| pLAR056 | $Ap^R$; Temperature sensitive; $P_{ENO}$-jK38 recombinase | (Riley et al., 2023) |
| pLAR058 | $Ap^R$; Temperature sensitive; $P_{ENO}$-R4 recombinase | (Riley et al., 2023) |
| pLAR060 | $Ap^R$; Temperature sensitive; $P_{ENO}$-TG1 recombinase | (Riley et al., 2023) |
| pLAR074 | $Ap^R$; Temperature sensitive; $P_{ENO}$-BXB1 recombinase | (Riley et al., 2023) |

| | | |
|---|---|---|
| pMTV210 | $Km^R$; oriR6K non-replicating integration plasmid with $P_{BAD}$ promoter (arabinose-inducible promoter) | This study |
| pMTV1127 | $Km^R$; oriR6K non-replicating integration plasmid with $P_{BAD::}$SYNPCC7002_A0358-BXB1 *attP* in pMTV210 | This study |
| pMTV1128 | $Km^R$; oriR6K non-replicating integration plasmid with $P_{BAD::}$SYNPCC7002_A0849-R4 *attP* in pMTV210 | This study |
| pMTV1130 | $Km^R$; oriR6K non-replicating integration plasmid with $P_{BAD::}$SYNPCC7002_A1188-SYNPCC7002_A1188-2-ϕK38 *attP* in pMTV210 | This study |
| pMTV1131 | $Km^R$; oriR6K non-replicating integration plasmid with $P_{BAD::}$SYNPCC7002_A2132-TG1 *attP* in pMTV210 | This study |
| pMTV1133 | $Km^R$; oriR6K non-replicating integration plasmid with $P_{BAD::}$SYNPCC7002_C0003-SYNPCC7002-C0004-BL3 *attP* in pMTV210 | This study |

**Supplementary Table 6:** Primers for Verification of *E. coli* Methylation Strains

| Primers | Sequence (5' to 3') | Uses and Notes |
|---|---|---|
| oMTV24 | GATCTCCTGTCATCTCACCTTG | With oMTV27 to confirm for Kanamycin marker in *E.coli* |
| oMTV27 | GAAGGCGATAGAAGGCGATGC | With oMTV24 to confirm for Kanamycin marker in *E.coli* |
| oMTV2222 | AGTGGCAGAACAGTGAAGGAAAC | With oMTV2223 to confirm SYNPCC7002_A0358 in *E.coli* |
| oMTV2223 | GAAATCCGAAAGACGTGCCATCAT | With oMTV2222 to confirm SYNPCC7002_A0358 in *E.coli* |
| oMTV2224 | GTTTGCTGGATGCGGAGGTATG | With oMTV2225 to confirm SYNPCC7002_A0849 in *E.coli* |
| oMTV2225 | CAGTGGCGGAATTGTTCGTGAAG | With oMTV2224 to confirm SYNPCC7002_A0849 in *E.coli* |
| oMTV2228 | CGTGAGCGTGTCTTCATTGTTGG | With oMTV2229 to confirm SYNPCC7002_A1188-SYNPCC7002_A1188-2 in *E.coli* |
| oMTV2229 | CTCGAACGGATGGATGAAGT | With oMTV2228 to confirm SYNPCC7002_A1188-SYNPCC7002_A1188-2 in *E.coli* |
| oMTV2230 | TACATCTTGGTCAGCGACTTTGC | With oMTV2231 to confirm SYNPCC7002_A2132 in *E.coli* |
| oMTV2231 | CGTGAAGCTCATCCAGAAACAA | With oMTV2230 to confirm SYNPCC7002_A2132 in *E.coli* |
| oMTV2234 | TTCTATCAACTGGACCCTGGTCG | With oMTV2235 to confirm SYNPCC7002_C0003-SYNPCC7002-C0004 in *E.coli* |
| oMTV2235 | CTGCAATAGGTGGTACTGGGAC | With oMTV2234 to confirm SYNPCC7002_C0003-SYNPCC7002-C0004 in *E.coli* |

**Supplementary Table 7**: Media A+ Recipe

| | **g $L^{-1}$** | **Final Concentration (mM)** |
|---|---|---|
| NaCl | 18 | 308 |
| $NaNO_3$ | 1 | 12 |
| KCl | 0.6 | 8.05 |
| $KH_2PO_4$ | 0.05 | 0.4 |
| $Na_2$ EDTA·$2H_2O$ | 0.03 | 0.081 |
| Tris Base + HCl (pH 8.2) | 1 | 8.25 |
| $MgSO_4 \cdot 7H_2O$ | 5 | 42 |
| $CaCl_2 \cdot 2H_2O$ | 0.352 | 2.39 |
| 1000X Trace Metals | 1 mL | |
| 1000X B12 | 1 mL | 0.000003 |

Solutions of $NaNO_3$, KCl, and $KH_2PO_4$ were prepared together at 50X concentration. Solutions of EDTA and Tris Buffer were prepared separately at 50X concentration and titrated to the appropriate pH with concentrated NaOH. The above solutions were then mixed with solid NaCl and 1000X Trace Metals and autoclaved. Separate solutions of $MgSO_4$ and $CaCl_2$ were made and autoclaved independently, then added to the solution once the media was cooled to prevent precipitation. B12 is made at 1000000X, filter sterilized, then diluted to 1000X for aliquots and used for media. Agar plates were prepared with 15g Bacto Agar in 500mL MilliQ $H_2O$ added to a 2X preparation of Media A+.

**Supplementary Table 8:** Trace Metals Recipe

| | **g $L^{-1}$ for 1000X Stock** |
|---|---|
| Ferric Ammonium Citrate | 3.81 |
| $H_3BO_3$ | 34.3 |
| $MnCl_2 \cdot 4H_2O$ | 4.3 |
| $ZnCl_2$ (2.3 uM) | 0.315 |
| $MoO_3$ | 0.03 |
| $CuSO_4 \cdot 5H_2O$ | 0.003 |
| $CoCl_2$(<1uM) | 0.006656 |
| $NiSO_4 \cdot 6H_2O$ (1 uM) | 0.2629 |
